# Limited stress and tissue-specific transcriptional and translational activity of transposable elements in mosquitoes

**DOI:** 10.1101/2024.02.15.580529

**Authors:** Elverson S Melo, Gabriel L Wallau

## Abstract

The mobilization of transposable elements (TEs) can either negatively affect the host’s fitness or contribute to the species evolution. TE protein expression is the first stage for transposition, but organisms developed defenses to control it. The intensity of regulatory mechanisms can vary among tissues, and in response to stress, it may facilitate TE activation across different species. Using hundreds of RNA-Seq and mass spectrometry experiments we calculated TE expression on twelve mosquito species. Most mosquito TE families exhibit constitutive RNA expression with abundant lncRNA production, yet only a limited number of proteins are effectively produced, in a tissue-specific manner. Under natural conditions, TEs exhibit distinct expression in somatic and germinal tissues, notably with pronounced repression in ovaries, associated with increased PIWI and AGO3 expression. Following exposure to abiotic stress and viral infection, certain TE families undergo altered expression. However, some stressors have no effects on TEs, or cause opposite effects in distinct species. Furthermore, repression predominates over induction in most cases. These data suggest that while some proteins are synthesized, the majority of TE transcripts function in a regulatory capacity. We also propose that the conventional notion of TEs being more expressed under stress conditions may not be universally valid.

**GRAPHICAL ABSTRACT:** 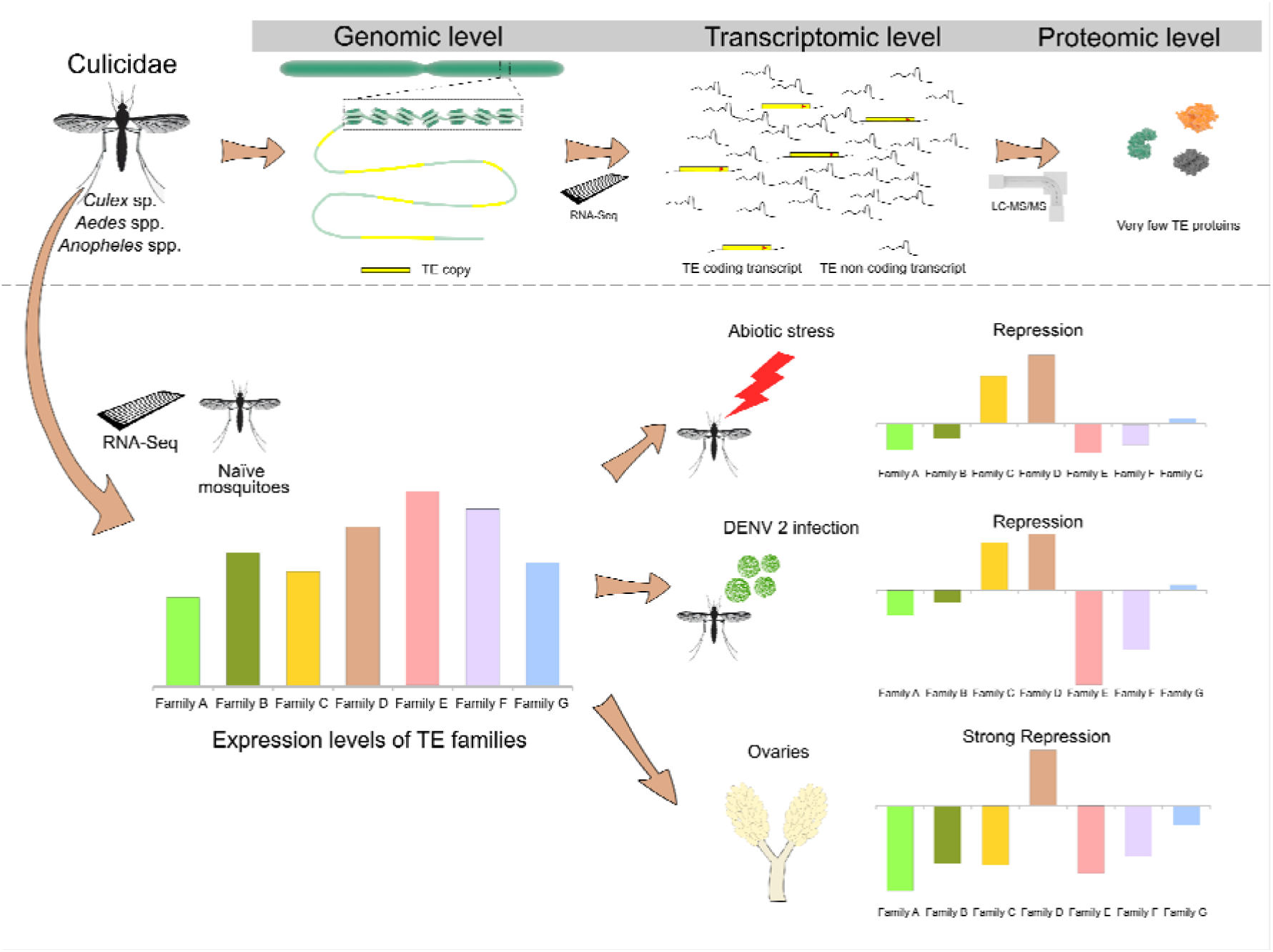

## INTRODUCTION

Transposable elements (TEs) are DNA sequences that mobilize between different loci of an organism’s genome and, during this process, generate new TE copies (1). Due to their selfish replicative nature, these elements attain high copy numbers constituting a large fraction of many eukaryotic genomes (2, 3). The majority of TE copies in a genome correspond to ancient transposition derivatives and hence most of the mobilome is composed of fragmented elements with limited transcriptional and translational activity (4, 5). However, the activity of the functional copies can generate new transposition events that may lead to genome instability and host fitness loss but occasionally contribute to adaptive insertions that increase host fitness (6). Various fitness-impacting insertions are known in eukaryotes such as: the somatic expression of a *mariner* element in *Drosophila melanogaster* inducing reduced larval crawling and viability of eggs (7); the expression of certain types of TEs, especially L1, is associated with several types of cancer in humans (8); the insertion of a hAT element into a gene produces an aberrant enzyme with negative consequences for lipid and protein biosynthesis in peas (9); the mobilization of different types of TEs that can lead to drug resistance in *Cryptococcus deneoformans* (10).

Due to the negative TE activity effects, molecular defense mechanisms that control TE expression at transcriptional and translational levels emerged and were naturally selected in eukaryotes, such as DNA methylation and several small-RNAs regulatory mechanisms (endo-siRNAs and piRNAs) (11, 12). TE repression by these mechanisms can also generate consequences to neighbor gene expression, such as the ERV repression effect leading to a decrease in the expression levels of dozens of flanking genes in human neuronal cells, which perturbates gene regulatory networks (13). Despite the occurrence of repressive marks that regulate TE RNA expression pre-transcriptionally, some copies are continuously transcribed generating abundant TE-derived mRNA that compose the whole organism’s transcriptome (14–17). These TE-derived transcripts show a wide range of structural diversity that varies according to the TE group and mobilization mechanism. Some transcripts represent the entire TE copy such as Class I TEs, that use an RNA intermediate during mobilization (18). Short TE transcript isoforms from Class I and Class II TEs can also be generated due to truncation or premature polyadenylation or splicing signals (19). Although some TE transcripts still retain coding capacity, the translation of these TE-derived transcripts is very difficult to detect, and very few studies propose to study it on a large scale (20).

Currently, the availability of large-scale transcriptomic datasets of some arthropod model organisms allows an in-depth assessment of TE activity at a genome-wide scale. Studies conducted mainly in vertebrates have shown that TEs display a tissue-specific transcriptional expression which has been attributed to specificities in histone modification as a result of differential expression of repressor proteins (21). In many vertebrates, the TE transcription level in germline and somatic tissues is substantial. Notably, the expression of TEs in testes generally exceeds that observed in other tissues, even considering that the expression of TE regulatory pathway genes is also higher in this tissue (15). The number of TE-derived transcripts can also be age-dependent, being more expressed in old flies of *D. melanogaster* (22) and modulated by viral infections in somatic tissues (23) or due to the stress response (24). There are a large number of studies that show the activation of some TEs in multiple species in the face of stress situations (25–29). Despite this, recent evidence shows a lack of general induction of TE expression in *Drosophila* strains subjected to stress (30). Despite highly heterogeneous results showing both activation and repression but also no transcription and translation modulation of TEs in natural and stress-related conditions, there is a widespread consensus in the scientific community that TEs are modulated by many stressors (31). The proportion of transcriptionally active TEs also varies substantially among species. For example, *D. melanogaster* TEs are much more expressed than human TEs, and it is expected that most TE families in that fly genome possess at least one active member (19).

Mosquitoes, compose a remarkably diverse group of insects encompassing around 3,500 species divided into two subfamilies, Anophelinae and Culicinae, both of which include mosquitoes of medical importance due to their capacity to transmit several viral and nematode pathogens, such as DENV and *Plasmodium* species to humans (32, 33). Comparative genomics of the genera *Culex*, *Aedes,* and *Anopheles* revealed many differences between Anophelinae and Culicinae, related to the gene copy number, sex determinants regions, and genome size (34). Previous studies have shown that transposable elements directly contribute to the genome size of species, varying from around 1% in some anophelines to more than 45% in some species from the *Aedes* genus (35). Despite the availability of several complete mosquito genomes and the large number of RNA sequencing experiments focused on mosquito differential expression of genes conducted until now, most studies on TEs concentrate on the investigation of small RNA pathways linked to TE silencing (36, 37). Only one study focuses on the discovery of transcriptionally active TEs in mosquitoes, this study, conducted in *An. funestus*, showed that several TEs of both Class I and Class II are transcribed, including a putative full-length element (38). Hence, for most species, the transcriptionally active TEs are still unknown, hindering the study of TE expression modulation and impact in response to external stimuli.

In this study, we investigate the transcriptional and translational activity of TEs in several mosquito species. Moreover, we leverage large RNAseq datasets designed to detect transcription-wide stress response in mosquitoes to evaluate a long-lasting question in TE biology: “Is genome-wide TE transcription and translational activity modulated by biotic and abiotic stressors?”. We show that many of the mosquito TE families are expressed but few transcripts encode a complete coding region. In addition, we also detected a consistent TE repression in mosquito ovaries, but no consistent mobilome-wide transcriptional repression or induction in testes or somatic tissue submitted to stress. Finally, at the proteome level, we observed an interesting tissue-specific pattern, where there are few TE proteins expressed in each tissue, and these proteins are generally not expressed in more than one tissue.

## MATERIALS AND METHODS

### Genome selection and TE annotation

We selected mosquito species with at least 10 public RNAseq runs available in the SRA database of NCBI: *Ae. aegypti*, *Ae. albopictus*, *Cu. quinquefasciatus*, *An. gambiae*, *An. coluzzii*, *An. arabiensis*, *An. merus*, *An. minimus, An. albimanus, An. funestus*, *An. dirus* and *An. stephensi*. For all those species, we previously characterized the total amount of transposable elements using *de novo* and homology-based approaches (35). Six out of twelve species have a better genome assembly available than the previous version used to characterize transposable elements in the publication. For these six species, we applied the TEdenovo pipeline (39) to uncover transposable elements, expecting to find less fragmented TE copies which will improve the functional analysis of the current manuscript.

Only *de novo* characterized TEs consensuses were used to annotate the mosquito genomes. We removed redundancy in the TE library of each species applying the 80-80 rule in step 6 of TEdenovo and filtering out repetitive elements without any TE features and chimeric TE consensus. After that, we mapped each TE consensus against the respective mosquito genome using blastn (BLAST 2.2.30+) using the following parameters: -task rmblastn, -dust no, -word_size 11, -soft_masking false. Only TE consensus with at least one match in the genomes was kept for the next step. Subsequently, we used the script filter-stage-1.prl of the RepeatScout software (40) to remove TE consensus that contained over 50% low-complexity regions (these consensuses were likely misassembled). The remaining final TE dataset was used to annotate the genomes of mosquitoes.

TEannot (39) pipeline was used to create TE annotation files (GFF format). Only Censor and BLR algorithms (step 2 of the pipeline) were used to map TEs to their respective mosquito genome, followed by the application of a statistical filter in step 3. We also use tblastx to annotate putative TE by homology using consensus present in the Repbase database (step 6 of TEannot). After these two TE annotation strategies the "long join procedure" was performed and only matches in the genome with a size higher than 300nt were kept in the gff3 annotation file. A custom script was used to remove any possible overlapping annotation of transposable elements (more than one TE in the same region) and to create clean GTF files (available on Figshare) used to estimate TE expression in the TEtranscripts package (41).

### Genome mapping

We searched all available RNAseq experiments in the SRA database for those twelve mosquito species until September 2020. We selected only RNAseq experiments that were conducted in the Illumina^®^ platform using paired-end reads (**Supplementary Figure 1** shows the number of RNA-Seq runs by species and by experimental groups). FastQC (http://www.bioinformatics.babraham.ac.uk/projects/fastqc/) and MultiQC (42) were used to evaluate the quality of all RNAseq runs. Trimmomatic (43) was used to trim any adapters and low-quality reads using the following parameters: LEADING:3, TRAILING:3, AVGQUAL:20, MAXINFO:75:0.5, MINLEN:35 (for a few samples, we varied the MAXINFO parameter, to adapt to read length). The STAR aligner (44), version 2.7.3a, was chosen to map RNAseq reads against mosquito genomes, as it easily allows multi-mapping during alignment. For differential expression experiments, we keep a maximum of one hundred alignments per read (--outFilterMultimapNmax 100) to allow capture reads from different copies of the same TE consensus. MultiQC was used to evaluate the mapping quality. Supplementary File 1 shows the quality of reads after trimming and the quality of BAM files. Experiments in which more than 50% of the reads did not align with the reference genome were excluded from the study.

### Evaluating transposable element expression

The program TEcount (part of the TEtranscripts package) was used to estimate the number of reads that map to each gene or TE. This program receives the .bam files, a GTF file from genes, and a GTF file from TEs and it outputs a count table for each experiment. Count tables from all experiments from the same mosquito species were concatenated. Genes and TEs with very low count values through all experiments were removed. This procedure was performed to prevent noise (wrong read mapping) from overestimating the number of TEs expressed.

The program TEtranscripts was used to perform differential expression analysis. Only experiments involving expression in somatic versus germline tissues (45–47) (**Supplementary Table 1**), and exposure to biotic and abiotic stress (48–57) (**Supplementary Table 2**) were analyzed. To avoid noise in the analysis we set --minread parameter equal to 10 and considered differentially expressed TEs only those with adjusted log fold change > |1|.

### Reconstructing transcripts of expressed TEs

To recover any potential transcripts derived from transposable elements, the BAM files of each species were submitted to Trinity (version: Trinity-v2.12.0) (58). The genome-guided Trinity Assembly was chosen to reduce the chance of assembling artifacts, as occurred in studies that used a Trinity assembly without any reference (59). In this mode, assembly is performed for reads that closely map each other in the genome, this prevents reads from similar copies of the same TE, scattered throughout the genome, from being assembled into a single copy. Another way to achieve more reliable reconstruction was to use read alignments from the same lineage of the reference genome for each species. To homogenize the discovery of TE-derived transcripts, only experiments conducted in adult mosquitoes were used for the assembly step. The evaluation of the assembled transcriptome completeness was performed using BUSCO (60) (version 3.1.0) with the Diptera database. TE-derived transcripts were uncovered by aligning the assembled transcripts to the TE consensus of each mosquito species (using blastn) and to the TE protein sequences of Repbase (using blastx).

TransDecoder (-m 200) was used to predict the Open Reading Frames (ORFs) for each TE-derived transcript to recover expressed potentially coding TEs. We include homology searches as ORF retention criteria of predictions, to avoid the estimation of spurious ORFs at the expense of the correct ORF of a TE, or even an ORF bearing a misassigned start site. A blastn mapping was performed against a database consisting of protein sequences of transposable elements available on Repbase (RepBase20.05 – 24195 proteins), plus TE protein sequences (3145 proteins) available on UniProt SwissProt (61). Additionally, we perform a search using HMMER 3.1 (62) to identify protein domains inside the ORFs, using the database ProfilesBankForREPET_Pfam27.0_GypsyDB.hmm (available at https://urgi.versailles.inra.fr/Tools/REPET), which includes Pfam (63) and GypsyDB (64) profiles.

Putative full-length TE transcripts were found using a custom script. To classify a transcript as complete, we evaluate if it had: I) all protein domains present in a given TE class, following the parameters determined by Wicker et al. (2007); II) all ORFs in the same strand; III) an untruncated start and end of each ORF present in the transcript.

### Discovery of TE-derived proteins

We conduct searches on the Proteomics Identifications Database (65) (PRIDE - https://www.ebi.ac.uk/pride/) to recover mass spectrometer experiments for all 12 studied species. Only three species, *An. gambiae*, *An. stephensi* and *Ae. aegypti*, showed satisfactory data (LC-MS/MS data, well-described experiments, adult mosquito as a sample). For *An. gambiae*, datasets evaluated were: Brain tissue - PXD000630 (66), whole body - PXD016300 (67); for *Ae. aegypti* we used: head tissues - PXD022665 (68), semen - PXD010293 (69), salivary gland - PXD002468 (70); finally, the dataset chosen to *An. stephensi* encompass multiple tissues of the whole mosquito body - PXD001128 (71). The .raw files of each experiment were downloaded from PRIDE and locally converted to .mzML format using ThermoRawFileParser (Peak Picking option set) (72) integrated into the SearchGUI interface (73).

For each species, a different search database was constructed encompassing 243 common contaminants, the TE protein dataset predicted by the TransDecoder from all TE copies of the reference genome and all TE-derived transcripts in each species, and the UniProt reference proteome of that species. To build the final dataset of putative TE proteins for each species (available on Figshare), firstly putative complete TE proteins were separated from truncated TE proteins. After that, the redundancy in both subsets was removed using the CD-HIT program (parameters: -c 0.98 -G 1) (74), this keeps only one isoform for each TE. Subsequently, the proteins in a subset of truncated proteins with high similarity (>99%) to proteins in the complete subset were removed using cd-hit-2d. Finally, both subsets are joined to build the TE protein dataset.

Proteomic analyses were conducted in SearchGUI, testing different search engines for each experiment: X! TANDEM (75), MS Amanda (76), MS-GF+ (77), Comet (78), and Tide (79). We also tested a *de novo* protein identification using Novor (80). Searches were performed with tryptic cleavage specificity; Enzyme: Trypsin; Precursor charge: 2-4; and peptide mass tolerance, fragment ion mass tolerance, the maximum number of missed cleavages, fixed and variable modification according to the methods described in the original manuscripts of the experiments. A Target-Decoy approach was used and a protein FDR (false discovery rate) of 0.01 was set as the cutoff for identifications. Results of different search engines were combined using PeptideShaker (81). A specific combination of search engines was chosen among those that increase the number of protein groups and TE proteins identified and minimize the PEP resolution.

## RESULTS

### Most mosquito TE families are expressed

TE sequences of 24 species of mosquitoes were previously genome-wide characterized (35), here we improved TE characterization to 12 out of those 24 species, including the newest high-quality versions for 6 mosquito genomes (**Supplementary Table 3**), and generated TE annotation files (available on Figshare) required for transcriptional expression assessment. Experiments from a total of 1060 publicly available RNA-Seq runs of many different conditions (**Supplementary File 2**) were mapped to their respective species’ genome. The quantification of mapped reads revealed that all 12 species express some TE family at the mRNA level. There is a high proportion of expressed TE families under different conditions varying according to species (51% to 94% of all TE families (Table 1)). This proportion is smallest in *An. dirus* (51%) and highest in *An. funestus* (94%).

**Table 1.**
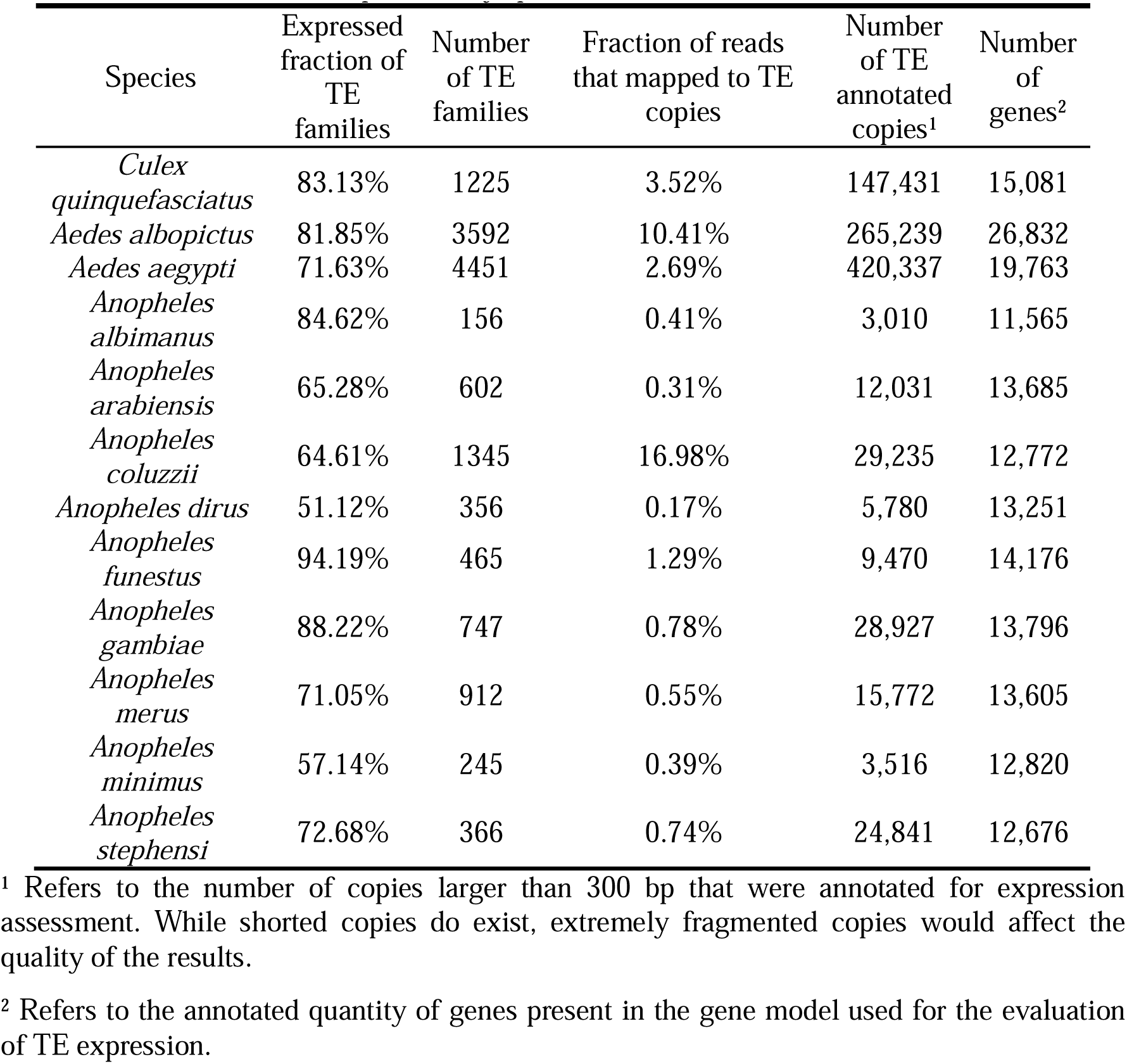
Profile of TE expression by species.

A commonly observed pattern from RNA-Seq data is a smaller average number of reads (normalized by Counts Per Million - CPM) mapping to TE families compared to the normalized average number of reads that map to genes for most mosquito species, except *Ae. albopictus* and *An. funestus* (**Supplementary Figure 2**). In other words, a TE family is typically expressed at lower levels than a mosquito gene, although some TE families show higher expression levels compared to many mosquito genes even considering that TE families are composed of several copies while most mosquito genes have a single copy. Therefore, the expressed mobilome constitutes only a minor fraction of the total transcript amount, on average only 3.19% of whole mapped reads in mosquitoes. However, this proportion varies depending on the mosquito species (Table 1) and no relationship was found between the fraction of reads that mapped to TE copies and the number of TE families (R=0.42, *p*=0.18) or the number of TE copies (R=0.25, *p*=0,43). This suggests that other factors, such as the age of the elements, the level of copy integrity (with the preservation of promoters), and the particularities of the arms race between transposable elements (TEs) and host silencing mechanisms, are also critical for the proportion of the transcriptome derived from TEs.

Given that most families of transposable elements are expressed in mosquitoes, a remarkable similarity is observed between the expression profile of the superfamilies (**Fig 1**) and the occurrence profile of these TE superfamilies in mosquitoes (**Supplementary Figure 3**). Thus, there are a few cases of TE superfamilies with detectable transcription. However, the expression level of superfamilies does not reflect their copy number in the genome. Expression also differs significantly between the different superfamilies (Kruskal-Wallis, p < 2.6 x 10^-16^), with Class I elements generally being more expressed than Class II elements. Among the most expressed superfamilies are Gypsy, Bel-Pao, I, and Jockey (**Supplementary Figure 4**).

**Figure 1.**
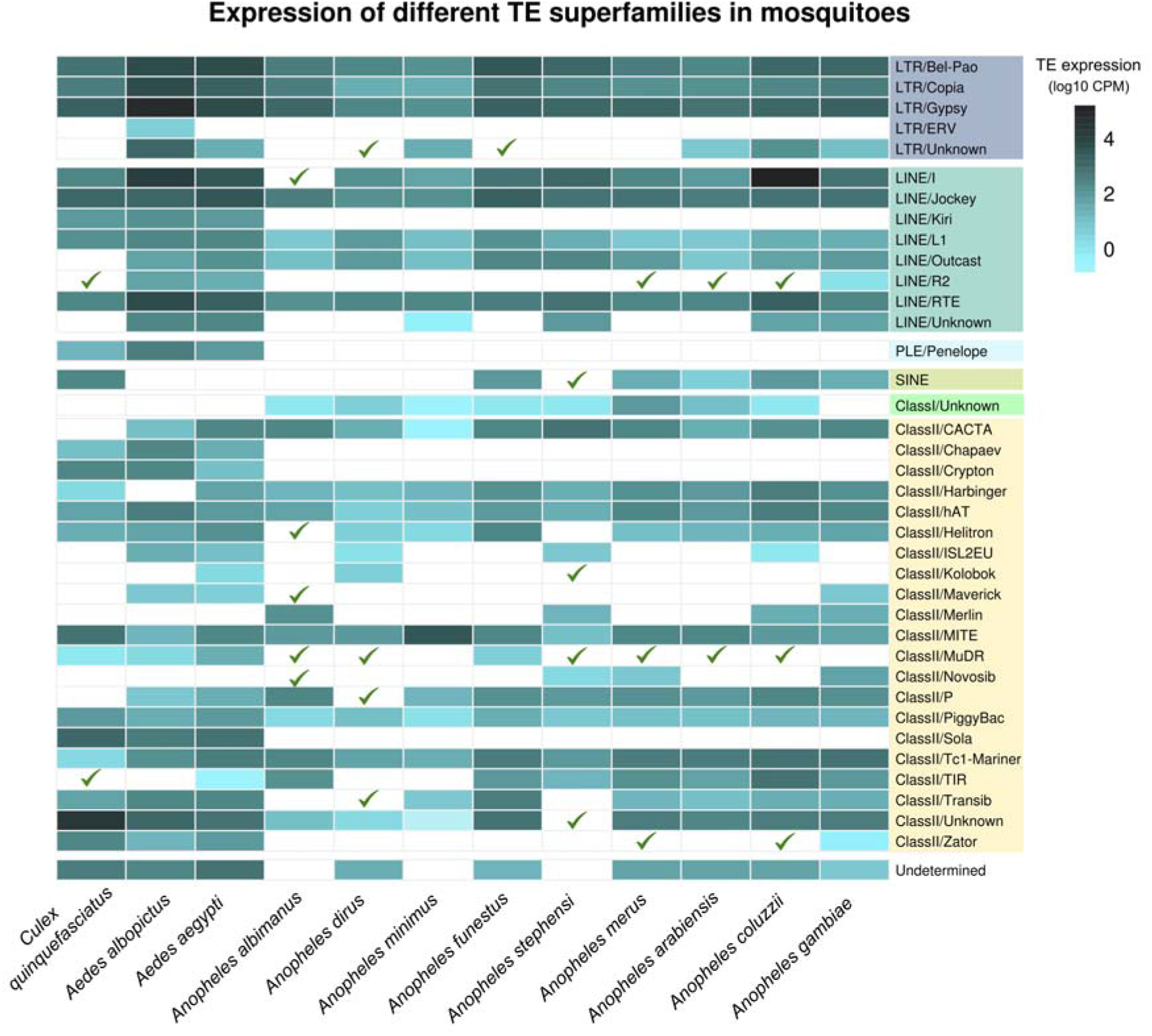
Heatmap of expression of different TE superfamilies in mosquitoes. The expression pattern of superfamilies closely mirrors their occurrence pattern in each species, reflecting the prevalent expression of the majority of transposable element (TE) families. The expression level of TEs reaches its lower threshold in *An. dirus*, which has only about 1650 reads mapping into TE copies for every million RNA-Seq reads mapped. The green checkmark icon in the figure stands for the occurrence of a superfamily that is not expressed in a specified species. The empty rectangles indicate the absence of a TE superfamily in a specified species.

### Mosquito TEs are strongly repressed in the ovaries

Ovaries, testes, and mosquito carcasses lacking reproductive tissues are quite common samples among mosquito RNA-Seq experiments available. To evaluate the TE expression between somatic and gametic tissues, we performed a differential expression analysis for testes, in seven species, and ovaries, in five species.

Transposable elements are much less expressed in the ovaries than in carcasses, although there are some cases of induction of TE expression in the latter. In 3 out of 5 species (*An. albimanus*, *An. arabiensis*, *An. gambiae*), more than a third of all expressed TE families are repressed in the female germline tissue, with particular emphasis on the ovaries of *An. gambiae*, where almost 50% of TEs were repressed (**Supplementary Figure 5** – Carcass vs ovaries). The repression pattern found in the ovaries is not observed in the testis. In this tissue, there are more repressed than induced TEs only for four species, for three other species, the inverse relationship is seen with a higher expression of TEs in testes than carcass (**Fig. 2 A** – Testes/Carcass). Regarding the TE type distribution, both Class I (LTRs and non-LTRs) and Class II are differentially expressed in ovaries and testes of mosquitoes, but with a prevalence of Class I TEs, mainly from Gypsy and Bel-Pao superfamilies (Fig. 2B).

**Figure 2.**
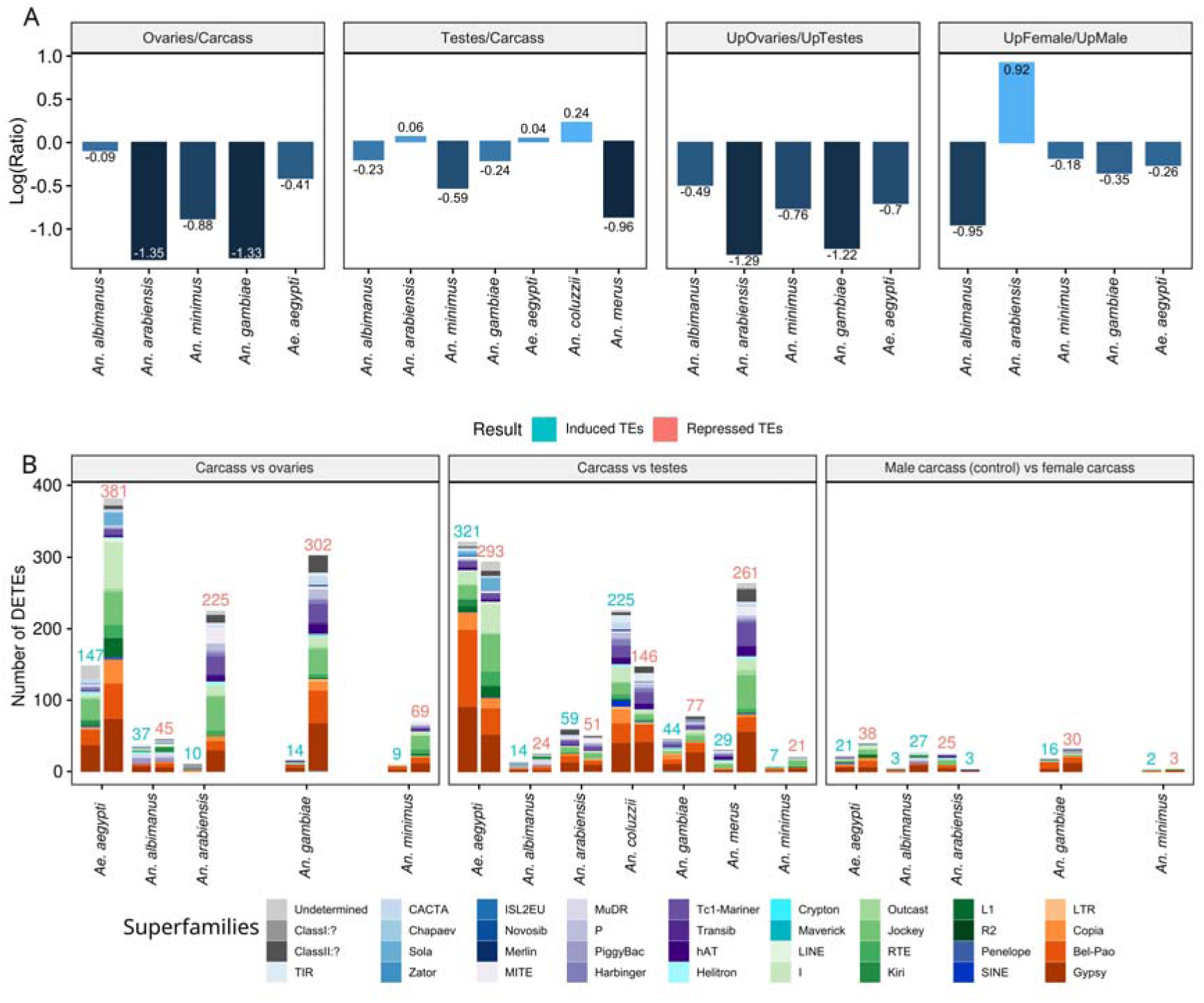
Differential expression of TEs in gonads and carcasses of males and females. Panel A shows the ratio between the number of upregulated TE families by the number of downregulated ones. In general, strong repression of TE expression is observed in the ovaries, but not in the testis. When compared between these two tissues, the number of TE induced in the testis is much higher than in the ovaries. Panel B considers the absolute number of differentially expressed transposable elements from each TE superfamily. The carcass was defined as a reference (control) for the carcass x ovaries and carcass x testis evaluation, while the male carcass was used as a reference for the comparison of male and female carcasses. For each species, the right column stands for induced TEs (blue numbers), and the left column repressed ones (pink numbers).

In general, more than 30% of expressed TE families keep the same level of expression in ovary and somatic tissues. This number is slightly higher, at least 70%, between testicles and carcass. However, for all species, there are more repressed than induced TEs in ovaries compared to testes, suggesting a strong repression of transposable elements in the ovaries of mosquitoes (**Fig. 2A**). In fact, TEs in ovaries show a completely different pattern of expression compared to the expression in somatic and male reproductive tissues in *An. albimanus* (Fig. 3A), *An. arabiensis* (Fig. 3B), *An. gambiae* (Fig. 3C) and *An. minimus* (Fig. 3D). In *An. minimus*, 44% of differentially expressed TEs (DETEs) (56 TE families) show a level of expression that differentiates from the level encountered in the carcass or the male germline (**Supplementary Figure 6**), this proportion is even higher in *An. arabiensis* (50%), *An. minimus* (58%) and *An. gambiae* (53%), and the overwhelming majority of these TEs are repressed.

**Figure 3.**
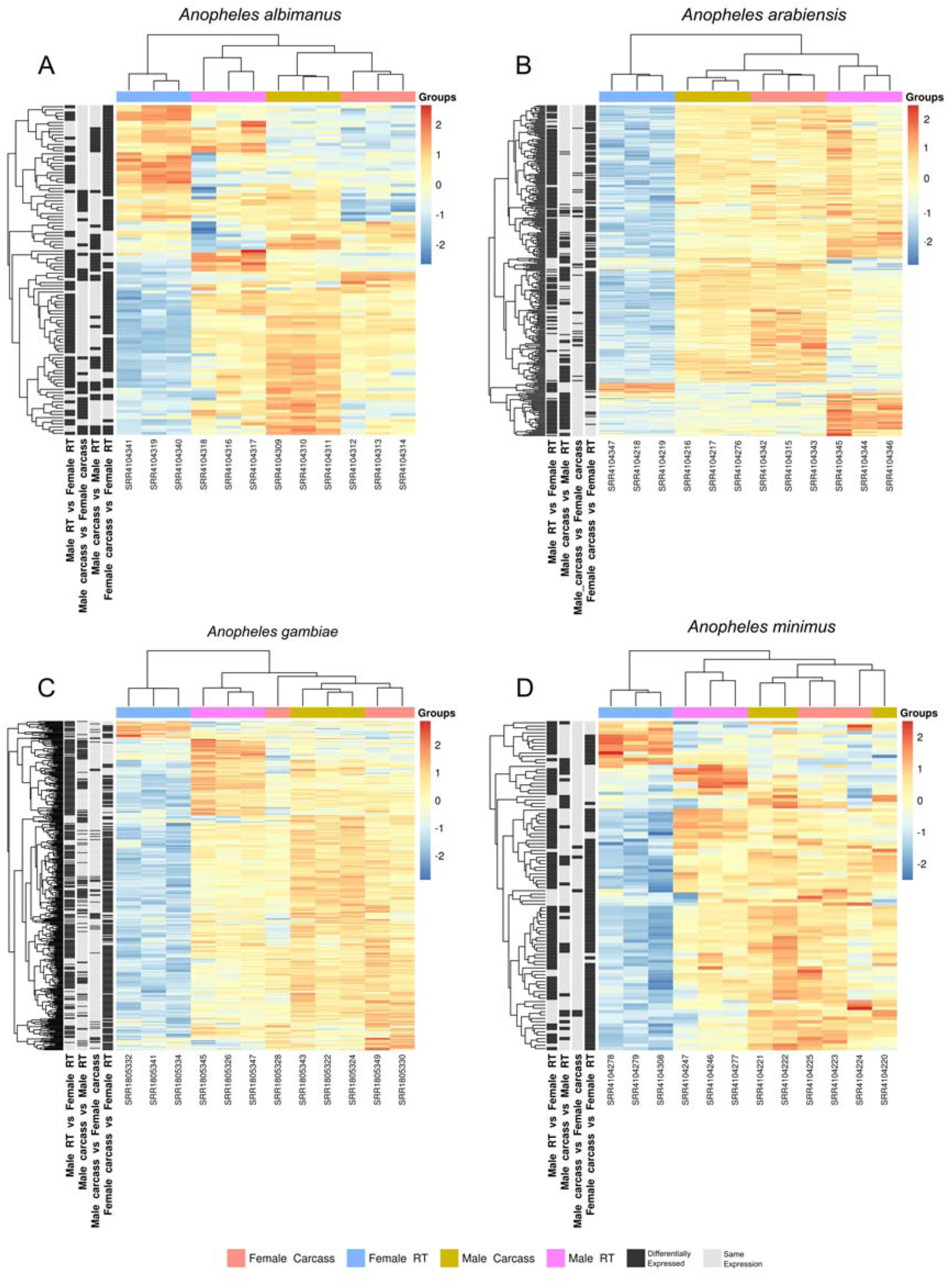
Expression pattern of different TE families in somatic and germline tissues. The heatmaps show the expression level of TE families (lines), in different samples (colored columns) of *An. albimanus* (A), *An. arabiensis* (B), *An. gambiae* (C), *An. minimus* (D). The four columns on the left, with cells in two shades of gray, indicate transposable elements whose expression differs significantly between two conditions, according to the DEseq2 results. RT means reproductive tissues.

The expression pattern of TEs in the germline tissues of *Ae. aegypti* was different from that observed in anophelines, only 25% of DETEs showed a different expression pattern in ovaries compared to the somatic and the male reproductive tissues. The germinal tissues of the male and female of this species have a TE expression pattern that is more similar to each other than that expression pattern observed in the somatic tissues, but with more TEs induced in testes than in ovaries (**Supplementary Figure 7**)

The sex influence in mosquito’s TE expression plays a pivotal role solely on gonads. In most mosquito species few TE families (<7%) present a somatic sex-biased expression (**Supplementary Figure 5**), except by *An. albimanus*, in which 22.8% of TEs are differentially expressed, 20.5% of them are less expressed in females. Interestingly, among this group of species, *An. albimanus* is the one that exhibits the highest prevalence of sex-biased gene expression, more than 15% of annotated genes (45).

To investigate potential causes related to this strong TE repression in the ovaries of species, we quantified the mRNA expression of genes encoding PIWI pathway proteins. There is an overexpression of most of the pathway proteins (Aubergine, Argonaute3, and PIWI) in the ovaries of *Anopheles* species compared to somatic tissues (**Supplementary Figure 8**). The same set of proteins that is overexpressed in the ovaries also shows changes in its expression in the testes; however, the expression in the ovaries is even higher (until three-fold higher). In *Aedes aegypti*, where there has been a great expansion of the gene family involved with the PIWI-piRNA pathway (82), 6 genes are overexpressed in the ovaries in relation to the female’s somatic tissues: AGO3, PIWI2, PIWI3, PIWI4, PIWI5, PIWI6 (**Supplementary Figure 8** – top panels). The relationship between the expression of these genes in ovaries and testes, nonetheless, demonstrates a more equitable distribution when contrasted with *Anopheles* mosquitoes: PIWI6 and PIWI4 are more expressed in the ovaries, AGO2 and PIWI2 are more expressed in the testes, while there is no change in the expression of the other genes.

The expression of most PIWI pathway genes does not vary between female and male somatic tissues of mosquitoes, in agreement with the small variation in the TE expression. Given that the expression level of certain proteins in the PIWI pathway changes when there is a shift in TE expression and stays relatively unchanged when TE expression differences are small, it is suggestive that the activation of PIWI pathway proteins is intrinsically linked to the expression of TEs in these species.

### TE expression is not strongly influenced by stress conditions

We evaluated four basic types of stress among mosquitoes: population density stress, when mosquito larvae were raised at a higher density per unit of space than usual; diapause stress, when the eggs of mosquitoes are placed under diapause-inducing conditions, such as lower temperature; chemical compounds stress; and salinity exposure stress. In all kinds of abiotic stress, except by *An. coluzzii* exposed to salinity, LTR retrotransposons made up most of the differentially expressed TEs (**Supplementary Figure 9**).

Interestingly, exposure to chemical compounds, such as ivermectin and deltamethrin in *An. gambiae* and *An. albimanus* respectively, practically did not change the TE expression (Fig 4A). Only a slight repression of the expression of 14 families (2.1% of the total) was observed in 6-day-old *An. gambiae* after ingestion of ivermectin. Exposure to diapause-inducing conditions was also unable to change the expression of most TEs in *Ae. albopictus* eggs, only 0.2% of TE families had their expression influenced by diapause conditions. In adult *Ae. albopictus*, this amount increases smoothly to 61 differentially expressed families, which corresponds to 1.8% of the total families of TEs of this species.

**Figure 4.**
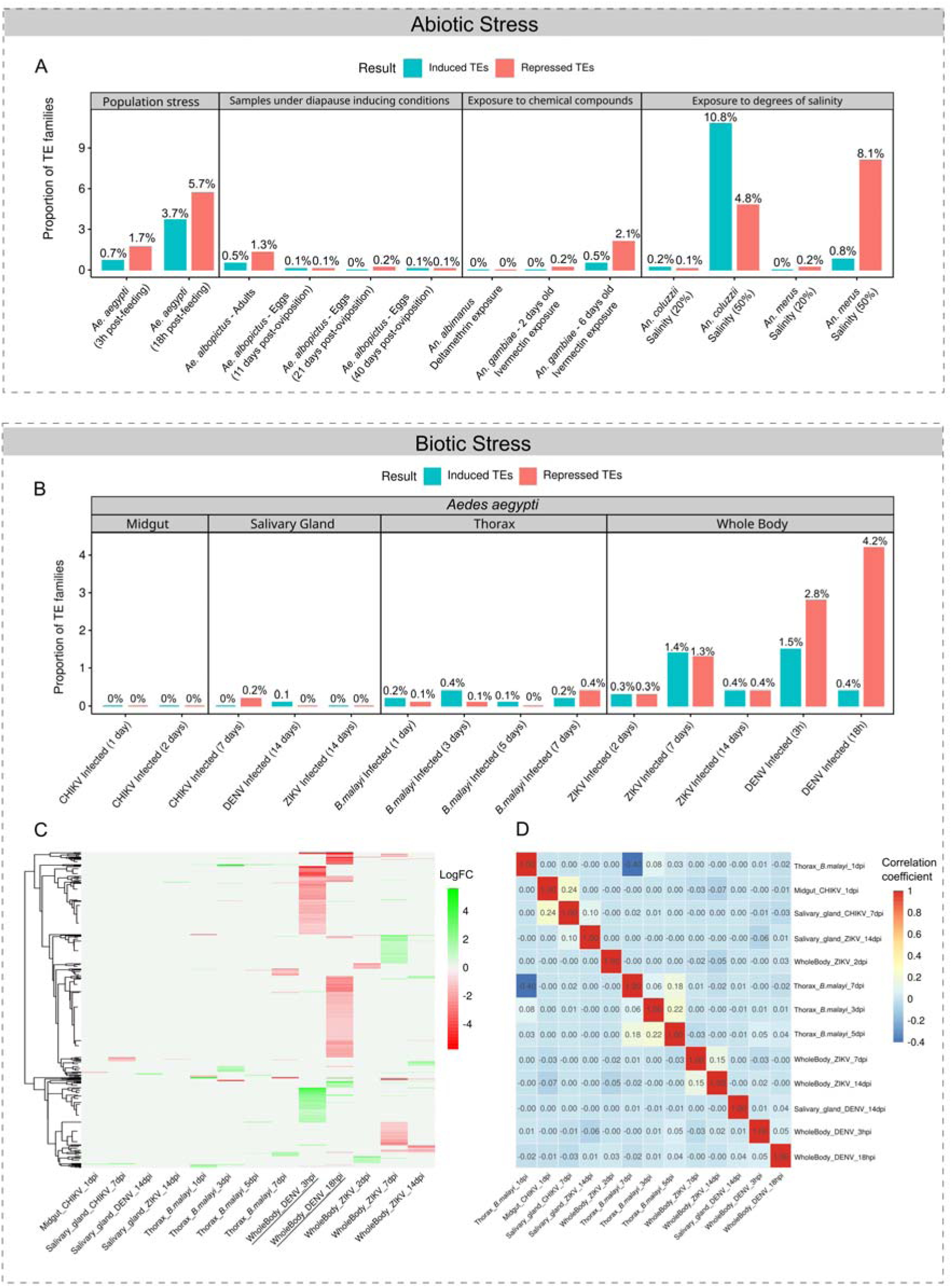
Differential expression of TEs in mosquitoes submitted to different stressful situations. (A) This panel shows the proportion of TE families for each species that are induced or repressed after abiotic stress and compared to a mosquito not exposed to stressful conditions. From left to right: I) “*Ae aegypti*” refers to females at different time points after blood feeding and derived from larvae grown under high population density (54); II) “*Ae. albopictus* – Adults” refers to females subjected to diapause-inducing conditions for 11 days (48); III) “*Ae. albopictus* – eggs” refers to eggs derived from parental females subjected to diapause-inducing conditions (49); IV) “*An. albimanus*” refers to mosquitoes that survived exposure to deltamethrin (50); V) “*An. gambiae*” refers to mosquitoes with 2 and 6 days old exposed to ivermectin (51); VI) “*An. coluzzii*” and “*An. merus*” refers to mosquito larvae exposed to different degrees of salinity (52). (B) This panel shows differential expression of TEs in *Aedes aegypti* after biotic stress caused by different pathogen infections, almost no significant difference in expression was observed in the midgut and salivary glands. (C) A heatmap showing the fold change in expression of different TE families (rows) that are differentially expressed in at least one condition. Each column stands for a different experiment and logFC values are calculated based on TE expression levels of their respective control group. (D) A correlation matrix of different samples based on the expression levels of DETEs.

Unlike previous stress conditions, exposure to salinity stress was able to change the expression of a significant fraction of transposable element families in mosquitoes (**Fig 4A**, right panel). In *An. merus* around 9% of TE families respond to salinity changes in the medium, while in *An. coluzzii* this value rises to 15.5% of the total TEs. However, this follows a stress intensity-dependent variation. At low salinity conditions, a small number of TE families have their expression altered, this changes considerably when the species is exposed to a salinity degree of 50%. In *An. merus*, a more salinity-tolerant species, most of the differentially expressed TEs represent families whose expression is repressed. Conversely, in *An. coluzzii*, a species that does not tolerate salinity, most of the differentially expressed TEs represent families whose expression was induced, around 11% of the TE families.

Population stress also altered the expression of TEs. In this case, more TEs are repressed than induced in *Ae. aegypti* (Fig. 4 A, left panel) whose larvae grew under high population density. In this case, a second stress factor was evaluated, the blood supply. Although it is not a type of stress that affects mosquito survival, feeding directly interferes with the physiology of the engorged female (83, 84). We show here that it also directly interferes with the expression of TEs, since the proportion of DETEs raised from 2.3%, after 3 h, to 9.4% of total TEs, 18h after blood feeding. In these two time points, the proportion of repressed TEs outweighed the number of TEs whose expression was induced by stress.

### TE expression in *Ae. aegypti* is affected by DENV2 infection

The considerable number of public RNA-Seq experiments of *Ae. aegypti* mosquito allowed us to verify the expression profile of TEs under different infection conditions and in different mosquito’s tissues (midgut, salivary gland, thorax, and whole body) (**Fig. 4B**). Infection with the DENV2 causes an expression change of a fraction of TEs families in the first hours after infection. 184 and 196 families of TEs showed some level expression variation at 3 hours and 18 hours after infection, respectively, in the mosquito’s whole body (**Supplementary Figure 10**). In this case, the number of TE families whose expression is repressed increases with the time of infection. Whereas the opposite pattern is seen for TE families whose expression is induced. Most of the TEs differentially expressed in one of the time points did not have their expression changed in the other, indicating that different families of TEs are modulated in different periods after the DENV2 infection stimulus (Fig 4C).

Infection with ZIKV is also capable of altering the expression of TEs in *Ae. aegypti*, but at a much lower level than DENV2 infection. Only 23 families of TEs are differentially expressed two days after contact with the virus. This number rises to 112 differentially expressed families after 7 days of infection and drops to 34 after 14 days (**Fig 4B**). The increase observed at 7 dpi is likely associated with the period (6 dpi to 10 dpi) during which ZIKV attains the highest peak concentrations across various mosquito tissues, which is followed by a decline in viral titration in the saliva after 14 dpi (85). In all these three time points the fraction of induced TEs is almost equal to the number of repressed TEs. However, the transposable elements that respond in the first days of infection, are almost completely different from the TEs that respond to ZIKV at 7 and 14 dpi, which have an overlap of DETEs between them (**Fig 4C**). Despite the variation in the expression of some families, about 95% of *Ae. aegypti* TEs remain with their expression unchanged after infection with these two viral types.

Although there is a variation in the expression of TEs in the salivary gland 14 days post-infection with ZIKV (2 families), this number is much lower than that observed in the whole-body sample (34 families). The infection of DENV and CHIKV viruses in this same tissue also indicates that virus infection has an extremely limited capacity to induce significant variations in the expression levels of TE in the salivary gland. There is also practically no difference in the expression of TEs in the midgut between 1- and 2 days post-infection with CHIKV. Moreover, almost all TEs that have some expression variation in both salivary gland and midgut are LTR retrotransposons (**Supplementary Figure 10**). In addition to viral infection, the response against parasitic infection was also evaluated. We noted that the expression of more than 99% of the TEs stays unchanged during the time course of infection by *Brugia malayi* (**Fig 4B**).

Overall, most TE families have their expression altered in only one experimental condition (Fig 4C). For this reason, there is a low correlation between TEs expression in different experimental groups. Infection with *B. malayi* is the group where TEs respond more similarly between samples, although this similarity is low (Fig. 4D).

### Putative full-length TE transcripts are expressed in most mosquitoes investigated

To recover the full-length sequences of all expressed transposable elements among the 12 species investigated, we *de novo* assembled the transcriptomes of each species. All assembled transcriptomes show high completeness, less than 10% of core genes are missing. Moreover, the completeness of the assembled transcriptomes was remarkably similar to the reference transcriptome of each mosquito (Supplementary Figure 11). This high transcriptome completeness is a good proxy that we are analyzing the almost full breadth of mosquito active transcripts, including transposable element transcripts.

*De novo* assembly reconstructed a variable number of contigs varying from 26,679 Trinity genes (very closed related transcripts) in *An. minimus* to 323,166 in *Ae. albopictus*, which has a higher number of transcripts isoforms (**Supplementary Table 4**). The number of TE-derived transcripts is highly variable between species (Fig. 5A). The species of the Culicinae subfamily have most of their genome covered by TE, and thus, they presented much more transcripts derived from TEs than the mosquitoes of the Anophelinae subfamily. In *Ae. aegypti* and *Ae. albopictus*, more than half of the assembled transcripts are related to transposable elements, contrasting to TE-poor species such as *An. albimanus* and *An. minimus*. However, even in these TE-poor species, it is still possible to find some expressed TE families. Even though the number of TE-derived transcripts is quite high in some species, the fraction of these transcripts that still have coding regions (with ORFs above 200 aa) is small. Overall, it corresponds to less than 10% of the TE transcript isoforms of the species (**Fig. 5B**). Of these isoforms of TEs with coding potential, the number of potentially complete isoforms is even smaller (**Fig. 5C**). Among more than 500,000 transcripts derived from TEs from the 12 species, just over 300 isoforms still show features of full-length TEs.

**Figure 5.**
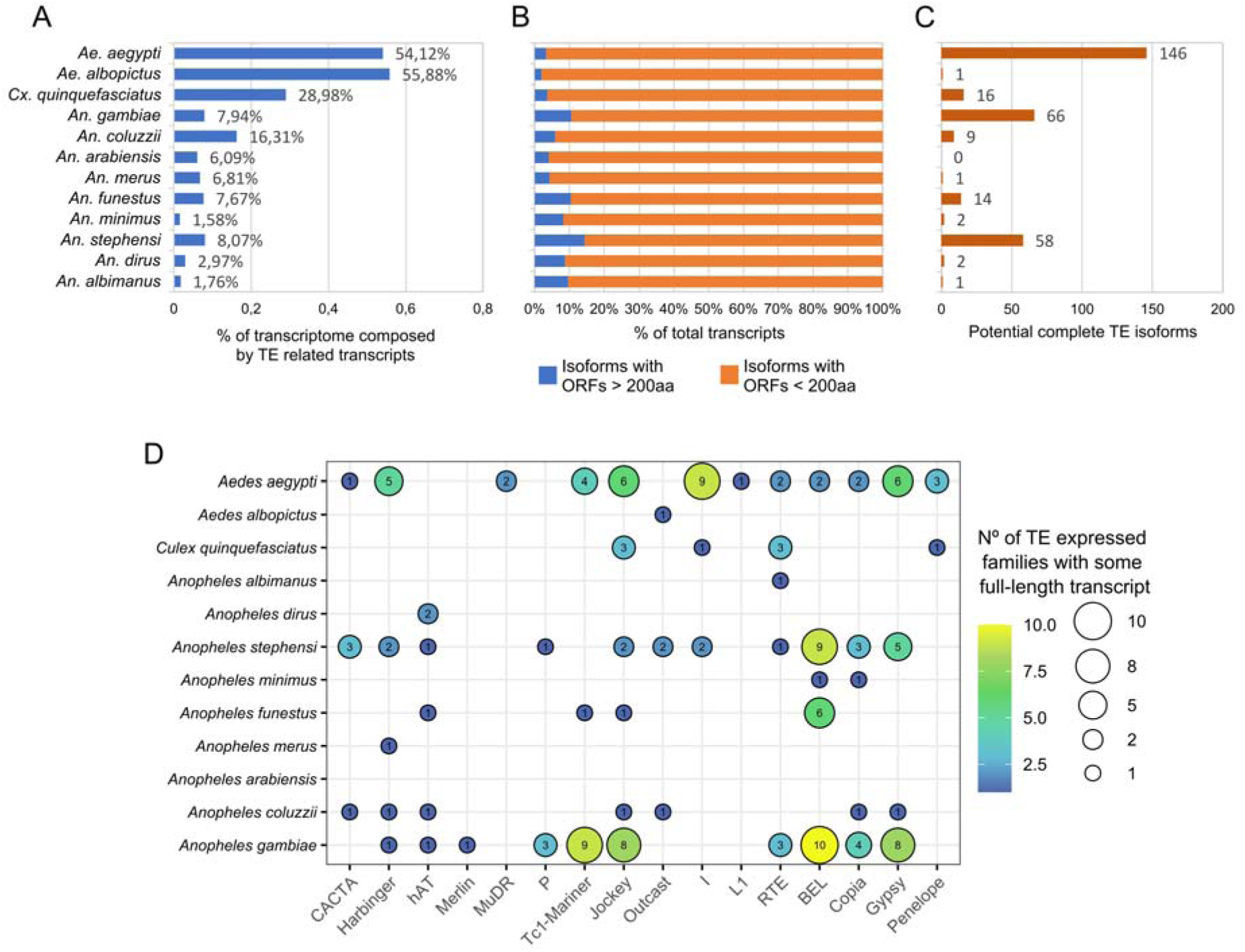
TE transcripts reconstructed by a *de novo* transcriptome assembly. (A) Fraction of the transcriptome generated from adult mosquitoes, consisting of transcripts associated with TEs. (B) Proportion of TE transcripts capable of generating significant ORFs. (C) The number of potential full-length TE isoforms among all recovered TE-related transcripts. (D) The colors and size of balloons are proportional to the number of families with complete isoforms within each TE superfamily (x-axis). The value inside the balloons stands for the amount of these families.

Out of this total, some isoforms are derived from the same TE family. Thus, the number of TE families that are still capable of producing potentially complete RNAs is smaller than the total number of isoforms. We also noticed that in some species such as *Ae. albopictus* and *An. albimanus*, only one TE family can still generate a potentially intact transcript. Meanwhile, in *Ae. aegypti* and *An. gambiae* 43 and 48 families are still capable of generating potentially intact transcripts, respectively, the highest numbers among species (Fig. 5D). The most prevalent superfamilies are the Gypsy and Bel-Pao LTR retrotransposons and the non-LTR retrotransposon Jockey.

### Expression of TE-derived proteins follows a tissue-specific pattern

Transcribed TE mRNAs that escape the host defense mechanisms at the transcriptional levels can be translated and generate proteins needed for their mobilization. To verify the ability of mosquito TEs to effectively produce proteins, we evaluate several publicly available mass spectrometer experiments. Only three species have satisfactory datasets for this kind of analysis: *An. gambiae*, *An. stephensi* and *Ae. aegypti*. They represent two genera, and within the *Anopheles* genus, two distinct sub-clades. The analysis was conducted using different algorithms to maximize protein identification. The approach results in a high spectrum identification rate (37.38%), which ranges from 10.67% to 58.37% in the different 22 samples (**Supplementary Table 5**), and a small (0,65%) posterior error probability resolution (PEP resolution (86)).

TE-derived proteins are expressed in all three species evaluated (Fig. 6). In total, 101 expressed proteins (available on Figshare) from 94 different TE families were found, and the expression of LTR retrotransposons stood out—more than half (56.38%) of the proteins were encoded by LTR retrotransposons. Out of this total, 23 proteins were derived from the Gypsy superfamily, 19 proteins from Bel-Pao, and 11 proteins from Copia. Non-LTR elements were also abundantly expressed, corresponding to 37.23% of all proteins found. All five LINE superfamilies (using the Wicker et al. (18) classification scheme) were found to be expressed. Nonetheless, certain superfamilies exhibit a more prevalent expression compared to others, as in the case of I and Jockey, which have 12 and 11 proteins expressed respectively, in contrast to the R2 group, which has only one representative expressed protein. Unlike retrotransposons, Class II TEs showed limited expression at the protein level. Only six elements of this class are expressed, all of them from the TIR order. Besides, they are expressed only in both species of the *Anopheles* genus and belong to the Tc1-Mariner, CACTA, and hAT superfamilies. All identified TE-derived proteins are described in **Supplementary File 3**.

**Figure 6.**
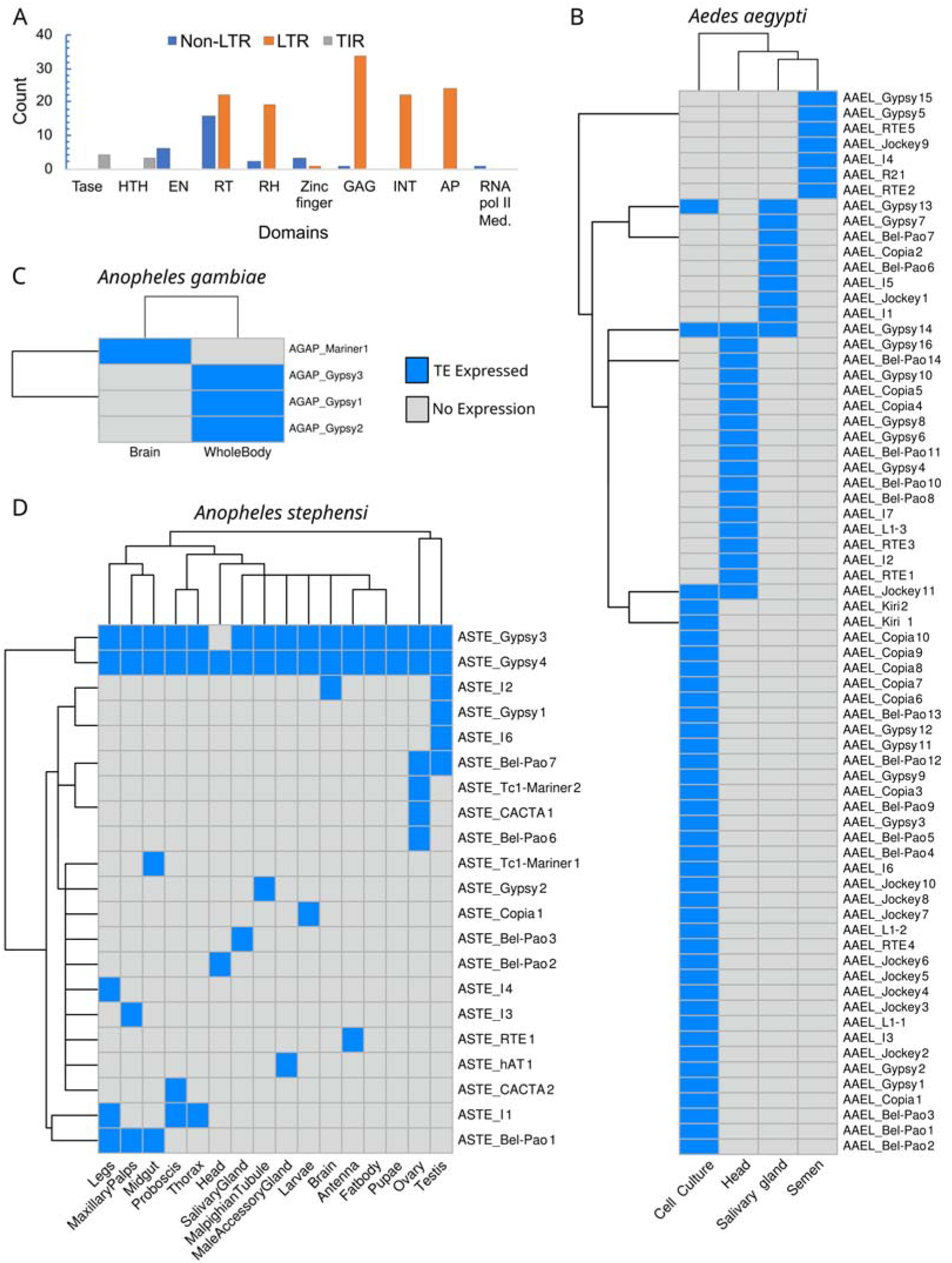
TE proteins identified by mass spectrometry. (A) Number of domains expressed in the TE proteins found in the 3 species, TASE = transposase, HTH = helix-turn-helix, EN = endonuclease, RT = reverse transcriptase, RH = RNase H, INT = integrase, AP = apurinic endonuclease, RNA pol II Med = RNA polymerase II mediator. Heatmaps showing TE proteins expressed in *Aedes aegypti* (B), *Anopheles gambiae* (C), and *Anopheles stephensi* (D).

Some of the identified proteins derive from incomplete TE copies, resulting in unequal distribution of protein domains (Fig. 6A). Within the LTR elements, the capsid GAG domain is the most often encountered, present in 34 of the 53 proteins of this order. This is the only domain present in the 2 TE proteins expressed in almost all tissues of the mosquito *An. stephensi* (Fig. 6D). Within the non-LTR elements, the distribution of domains exhibits an even more irregular pattern. Most of the non-LTR TE proteins only coded for the reverse transcriptase (RT) domain. Sixteen proteins without conserved domains were found, considering all protein groups analyzed. Despite this lack of domains, confirmation of the TE-derived origin of their ORFs has been obtained. Most of these proteins belong to non-LTR elements, which are known to have a second coding region (ORF1) whose function is not yet fully understood, but recent findings gathered evidence of their involvement in nucleic acid binding function and chaperone activity (87, 88). There are 25 complete proteins with canonical domains (18) characteristics of each superfamily, six in *An. stephensi*, four in *An. gambiae* and 15 in *Ae. aegypti*. Most of them represent GAG protein (a structural protein) (15 proteins), followed by fused Pol-Gag proteins (5 proteins) and stand-alone Pol protein (3 proteins) from LTR retrotransposons. Only two transposases, both from Tc1-Mariner elements, show features of conserved domain and complete ORF. These two Tc1-Mariner elements, one found in *An. gambiae* and the other in *An. stephensi*, are highly suggestive of TEs that probably still retain the ability for mobilization within these three mosquito species, as indicated by the completeness of the produced transposase at the protein level and its simple mechanism of action.

*Ae. aegypti*, the species with the largest fraction of genome covered by TE, is the one with the highest number of TE proteins expressed (69 proteins), even though there were only four tissues analyzed in this species (Fig. 6B). Despite this, more than a half of those proteins (36) is only expressed in cell culture of Aag2, an immortalized cell lineage that is frequently used in *in vitro* studies for investigation of immune response against pathogens (89). Although we investigated 16 different tissues in *An. stephensi*, only 21 TE expressed proteins were found (Fig. 6D), but in most cases, there are merely 3 or 4 proteins expressed in each specific tissue type. Finally, with only four proteins found in *An. gambiae* (Fig. 6C), although this small number is probably related to the amount of mass spectrometry available data, only for two tissues.

Interestingly, the expression pattern of TE-derived protein is highly tissue-specific. Only one (Gypsy_14) and four (Gypsy3, Gypsy4, Bel-Pao1, and I1) TEs expressed proteins in at least three different tissues of the species *Ae. aegypti* and *An. stephensi*, respectively. Most of these proteins present only the GAG domain, except the protein expressed by ASTE_I1, which is a truncated protein that encodes only the reverse transcriptase domain. Two of these proteins (Gypsy3, Gypsy4) are both expressed in almost all tissues of *An. stephensi*. Another interesting pattern found in this species is the presence of a quite different TE expression profile in the germinal tissues, which cluster together in the dendrogram (Fig. 6D). Both the ovary and testis express more proteins than any other tissue of *An. stephensi*, and seven out of nine proteins are exclusively expressed in these organs.

## DISCUSSION

Transposable elements evolve mostly neutrally after the invasion of a new host genome and hence most element copies are expected to accumulate mutations over time and degenerate at the functional level. The antiquated perspective deemed transposable elements (TEs) inherently harmful, which would be the cause of the suppression of genomic activity due to potential detrimental effects such as harmful mutations and genomic instability. However, contemporary research has unveiled a more nuanced understanding. While certain TEs may truly negatively affect their host genomes, others may play pivotal functions in evolution and genomic diversity, contributing significantly to host organism adaptation. The few full-length TEs and a much larger fraction of fragmentary elements scattered across the genome may still substantially influence the host gene expression (90). High TE expression at transcriptional and translational levels is expected to induce a heavy load to the host molecular machinery and when associated with TE mobilization may lead to detrimental effects on the host fitness, but most of these findings with precise TE-phenotype links are restricted to humans, mouse, and *Drosophila*. As an example, in humans, chimeric TE-gene transcripts are actively modulating oncogenesis (91). On the other hand, TE expression in non-model species is not well understood. In this study, we successfully constructed a comprehensive annotation of transposable element (TE) copies and made this annotation available to the scientific community. The analysis of these copies at the mRNA level revealed that most of the species’ mobilome is expressed.

### Abundance and structure of TE-derived transcripts

Here, we showed that most mosquito TE families are transcribed at the mRNA level and hence have at least one TE copy generating such RNAs. The high number of expressed TE families contrasts with the classical notion that most TEs are silenced and only a very limited number successfully achieve transcriptional expression (92). However, these findings are consistent with recent observations in other animal species, exemplified by the widespread TE expression observed in the salamander *Ranodon sibiricus* (93), and rats *Rattus norvegicus*, where 54% of TEs are expressed in at least one different organ (94). Furthermore, even in unrelated organisms such as the pathogenic fungus *Zymoseptoria tritici*, the majority of TE families are expressed in some strains under natural conditions (95). At the superfamily level, the most expressed TEs in mosquitoes are from the LTR order, followed by non-LTR ones. This corroborates studies conducted in different organisms such as plants, fungi, and vertebrates, which consistently demonstrate that the majority of TEs recovered in transcriptome assemblies are Class I elements (16, 17, 96). This pattern suggests that retrotransposons are more prone to transcribe in all eukaryotes, probably due to their mode of propagation. However, despite being considered non-autonomous TEs and lacking coding proteins, MITEs are also found expressed in all mosquito species evaluated, with particularly high expression in *An. minimus*, mirroring observations in *Z. tritici* lineages (95).

The level of the expressed mobilome in mosquitoes is similar to the expression levels observed in vertebrates, but to a lesser extent, accounting for 3.19% of the mapped reads, as opposed to the 6.55% observed in vertebrates (15). However, this proportion varies significantly among mosquito species, with less than 1% of the reads for some species and more than 10% in others. In mice, the TE expression is also much lower than gene expression and only 12-14% of RNA-Seq reads from embryonic cells map to TE copies (97). These pieces of evidence raise a hypothesis that for eukaryotes, there is a basal and ubiquitous expression of TEs.

Although expressed in low abundance in mosquitoes, the number of isoforms generated by TEs is quite variable between species, from ∼1,5% to ∼16% in Anophelinae mosquitoes to almost nearly 55% in both *Aedes* species. The TE transcripts proportion observed in Anophelinae mosquitoes transcriptome is more similar to the proportion observed in various species from different phyla: Mollusca, *Pomacea canaliculata* (1.9% of the transcripts) (98); Nematoda, *Heterodera avenae* (3% of the transcripts) (99); Chordata, *Rhinella marina* (∼8.2% of transcripts) (100) and *Corydoras maculifer* (4.68% of transcripts) (101); Basidiomycota, *Phakopsora pachyrhizi* (1.62% of the transcripts) (96). Although multiple isoforms of TE transcripts are produced, the difference between expression levels and transcript diversity in mosquitoes can also be quite high. As an example, in *Ae. aegypti*, TEs represent only 2.69% of all reads sequenced, but represent about 54% of assembled transcripts. This suggests that the difficulty/impossibility of annotating part of the transcripts in *de novo* transcriptome assembly studies is largely due to the presence of TE-derived transcripts. In fact, this large fraction of TEs in the transcriptome of *Ae. aegypti* is similar to the 55% fraction of unannotated contigs in the transcriptome (trinity-assembled) of *Ae. fluviatilis* (102). The use of only the NCBI nr database for TE discovery may have been responsible for the very low number of TEs found in this species (61 out of 58,013 assembled transcripts), as the TEs are underrepresented in the NCBI nr database. In addition to the prior recommendation to utilize TE databases during annotation to enhance the annotation of assembled transcripts (103), we propose the concurrent utilization of the Repbase protein database alongside the species-specific TE consensus database.

Despite TE transcripts reconstruction, a question remained open after the initial analysis of expression: do the transcripts originate from intact TE copies, or do they originate from defective TEs? Most of the assembled transcripts represent defective elements, characterized by small and truncated ORFs or with transcripts with no coding potential, similar to vertebrates, where most TE-derived transcripts are truncated (100, 104). Although *de novo* transcriptome assemblies suffer from incomplete transcripts, we used high-depth runs and genome-guided assembly and obtained high-quality transcriptomes. Therefore, the presence of TE transcripts without ORFs represents, in most cases, the occurrence of long non-coding RNAs (lncRNAs). In fact, as previously demonstrated, a considerable portion of the lncRNAs from *Ae. albopictus* and *Cx. quinquefasciatus* is associated with TEs (105). Classically, lncRNAs form a group of RNAs with heterogeneous regulatory functions that only share a length of more than 200 bp and lack coding regions (ORFs likely to generate polypeptides larger than 100 aa) (106). Despite the classic concept, a consensus proposal for the classification of non-coding RNAs suggests a subdivision into three groups. This would result in the different categorization of transcripts traditionally called lncRNAs, now divided into two subcategories: small Pol II transcripts (50–500 nt) and lncRNAs (>500 nt) (107). Despite the prevalence of small Pol II transcripts in mosquitoes, both categories of transcripts are represented in the transcriptome associated with TEs in mosquitoes (**Supplementary Figure 12**). In addition to the absence of ORFs, these transcripts associated with TEs also have other general characteristics of lncRNAs, such as low expression, which may be essential for their regulatory role, and the high number of isoforms (107), most found in Culicinae mosquitoes. This evidence suggests that the majority of TEs still expressed at the RNA level in mosquitoes have a regulatory role. Nevertheless, despite the low likelihood of TE mobilization in these species, new copies may be generated, as some transcripts still potentially code full TE proteins (Fig 5C).

### Transposable elements and proteomics

Until now, comprehensive studies assessing the large-scale production of TE-derived proteins have been exceptionally scarce. Six years after the first publication focusing on large-scale proteomics of transposable elements by Maringer et al., few large-scale proteomics studies have identified the proteins derived from TEs (20, 108, 109). The evaluation conducted in mosquitoes brought valuable contributions to the TE research field, making it clear that the expression profile of TEs at the protein level is highly tissue-dependent, and only a few TEs are expressed at the protein level. This low number of identified TE proteins was expected since the uncontrolled production of TE proteins can cause a deleterious impact on the host (22). However, the large difference between the high number of TE transcripts with ORFs and the small number of detected proteins may indicate post-transcriptional regulation of TEs—such as, for example, by piRNAs—rather than a process of transcriptional gene silencing, such as DNA methylation, given that the latter mechanism would hinder the abundant formation of transcripts (110, 111)

Some interesting insights derived from the observed TE protein expression pattern are: I) the use of cell cultures is not ideal for studying TE expression; II) differences in TE protein expression between tissues are more striking between somatic and germinal tissues. The tissue expression specificity and the different TE expression patterns observed between Aag2 cells and the other *Ae. aegypti* tissues, confirm that the TE expression is entirely distinct between cell culture lineages and mosquito tissues. Thus, studies that focus on assays with experimental conditions performed in cell culture will not adequately reflect the natural expression of transposable elements observed in mosquitoes. The tissue-specific expression pattern, with the formation of two clusters of TE expression, which differentiates somatic and germinal tissues in *An. stephensi*, reflects the remarkably differentiated profile of TE expression in the ovaries and testes, as evidenced by the several RNA-Seq assays. This difference in proteins is probably a result of the activation of different metabolic and regulatory pathways, as discussed in the next topic (15, 112).

Proteomic analysis indicated rare cases of TE protein domestications. Most of the identified ones still have TE-conserved domains, which contrasts with the TE protein domain degradation that normally occurs during the domestication phase of a TE protein into a new organism’s gene. Only two proteins were detected in *An. stephensi* deviate from the tissue-specific expression pattern and are expressed in almost all tissues. These proteins, XP_035912068.1 and XP_035912069.1, compose the predicted proteome of the species deposited at the NCBI, classified as “uncharacterized protein”, and still have a partially conserved GAG domain. Their orthologs are present in several species of the genera *Anopheles*, *Culex,* and *Aedes* and some other arthropods of the Insecta class. These two proteins represent putative cases of TE domestication in mosquitoes, but their function has not yet been deciphered.

From a methodological point of view, combining predicted ORFs from transcriptomic and genomic datasets increased mosquito TE protein identification. This suggests a new method of analysis compared to that previously recommended in the literature, which focused on the use of proteomics informed by transcriptomics (PIT) (20). On the one hand, one potential contributor to the observed disparities in results lies in the utilization of publicly accessible RNA-Seq and mosquito mass spectrometer data as the primary source for assessing TE expression, which represents heterogeneous samples. On the other hand, it was previously shown that, sometimes, the correlation between transcriptome and proteome is only modest (113, 114). Therefore, we suggest the use of PIT in conjunction with the ORFs of the TEs predicted directly from the genome of the species.

### Influence of sex on TE expression, with a focus on germinal tissues

The level of TE expression in different mosquito tissues and species varies substantially, but overall, there is a strong TE repression in ovaries. The repression of TEs in mosquito testes, when compared to somatic tissues, is not as pronounced as observed in the ovaries. This TE expression pattern in mosquito gonads adds another layer of evidence indicating a sex-biased TE expression in germline tissues of most animals, with overexpression of TEs in the testes. This pattern of expression is found in many species, such as the case of *D. melanogaster* (115), a species that belongs to the Diptera order (just like mosquitoes), and the case of very distantly related organisms, as vertebrates, where TEs show a higher expression in testes than in ovaries (15, 93). In certain species, such as *Rattus norvegicus*, TE expression levels in the testis surpass those in most somatic tissues (94). The repression of TEs in ovaries relative to somatic tissues and testes is nearly a rule among animals. TE expression in mosquitos is in line with high repression in the ovaries (between 8% and 46% of the TEs of a species) and that had been previously observed in other organisms (15, 93, 115–117). Only in a few species, such as the Japanese flounder *Paralichthys olivaceus* and two species of crow, TEs are more expressed in the ovaries than in the testes (118, 119). In the latter case, the expression in the ovaries is so high that it surpasses the expression of TEs in somatic tissues.

Regarding the types of TEs expressed, although the most common orders of TEs (LTRs, LINEs, and TIRs) are differentially expressed between somatic and germline tissues of mosquitoes, this behavior appears to be specific to each group of animals. In the grasshopper species *Locusta migratoria* and *Angaracris rhodopa*, only the LTR Gypsy and Copia are differentially expressed between these two kinds of tissues (120).

If a transposition event occurs in somatic tissues, it can cause damage to the organism or can be neutral. However, this new genomic organization is not inherited through generations because vertical transmission of somatic cells does not occur in animals. To propagate and colonize genomes, transposable elements need to express their proteins in the tissues responsible for gametic production. This leads to an arms race between TEs and hosts. If on one hand, TEs must express a substantial number of proteins essential for transposition, increasing the number of their genomic copies. On the other hand, the host suppresses the TE expressions to minimize their potential deleterious effects. To keep genome integrity, different organisms developed different defense systems, which can be broadly classified into repressor proteins and piRNAs-mediated silence. While the first mechanism acts only in vertebrates, the latter occurs in virtually all eucaryotes and can act in the host’s nucleus and cytoplasm (117). One of the causes of this more accentuated repression in the ovaries may be a greater production of small RNAs in this tissue, mainly piRNAs. In many arthropod species, piRNA molecules are strongly expressed in germinal tissues and to a lesser extent in somatic tissues, with somatic expression being an ancestral trait of arthropods. A significant fraction of the piRNAs present in the germinal tissues of these animals engage in the control of TEs. Even in cases of somatic piRNAs, a fraction of these RNAs still map to transposable elements, although most piRNAs appear to silence genes and in some cases, viruses (121). The piRNA involvement in the TE repression in ovaries, as in *D. melanogaster*, has been previously demonstrated. In this species, ovaries exhibit a higher abundance of piRNAs compared to testes, in addition to having an overexpression of piRNA pathway genes (122). The same pathway seems to be active in vertebrates, such as the zebrafish, in which piRNAs are expressed only in germline tissues and, among those piRNAs that map against TEs, in higher abundance in the ovaries than in the testes (123). However, this is not always the case, in salamander *R. sibiricus*, there are more piRNAs in the testis than in the ovaries (93). Although there are currently no RNA-Seq experiments targeting small RNAs to confirm the hypothesis of a higher amount of piRNAs acting in the mosquito ovaries, we can use expression data of proteins that participate in some steps of the silencing cascade.

Although the knowledge of the PIWI pathway in mosquitoes is not so extensive as in the *Drosophila* genus, many different PIWI proteins have been identified in these organisms (82). Mosquitoes have a variable number of proteins involved in the pathway, with a high expansion of PIWI proteins in the *Aedes* genus. For example, the PIWI5, PIWI 6 and PIWI7 transcripts present a high similarity with only one Aubergine-like transcripts of *Anopheles* mosquitoes (124). Our results showed that mosquito PIWI transcripts are overexpressed in ovaries compared to adult somatic tissues. These findings align with prior observations in *Ae. albopictus*, where PIWI2, PIWI3, PIWI6, and AGO3 exhibit high expression and PIWI7 is downregulated in ovaries (124), the same pattern observed in our results with *Ae. aegypti*. Although the piRNA pathway in *Aedes* mosquitoes is generally associated with antiviral defense and control of EVEs, this seems to be an additional function once the PIWI proteins are essential for the proper functioning of the TE silence pathway in most animals (82). The absence of PIWI proteins directly impacts TE in both vertebrates and invertebrates, for example, the knockdown of the Piwi2 protein in the shrimp *Penaeus monodon* caused the overexpression of some transposable elements and a decrease in sperm production (125). Additionally, in the Japanese flounder *P. olivaceus*, TE expression becomes more intense in the ovary than in the testis when the piRNA pathway proteins are not expressed in the ovaries (118). Our hypothesis posits a direct relationship between the robust TE repression observed in mosquito ovaries and the overexpression of specific Piwi and AGO3 transcripts. On the contrary, the downregulation or lack of differentiation in AGO2 expression (associated with the siRNA pathway (126)) within mosquito gonads provides stronger evidence in favor of the hypothesis that piRNAs, rather than siRNAs, actively contribute to TE regulation in mosquito germinal tissues.

Despite the significant impact of sex on the expression of TEs in germinal tissues, gender has minimal influence on the TE somatic expression of most evaluated species, except by *An. albimanus*. A small effect of sex was also observed in the studies (94, 116, 127), but in some cases, there is a strong somatic sex bias, mostly associated with TEs in sexual chromosomes (119).

### Stress response and global mobilome expression: an intriguing question

There are other sources of TE expression variation in addition to the effect of regulatory pathways between somatic and germline tissues. In some organisms, the somatic TE expression varies depending on the age (22), being highly expressed in elder specimens (128), or it can vary in response to many kinds of stress, like change in temperature, excess pollution, exposure to toxic compounds and radiation, and even psychological stress (24, 129). TE expression in stressful situations is still a matter of debate. Many different studies argue that there is an increase in TE expression in stressful situations. However, many of these studies focus only on a few types of TEs in situations such as aging stress (130), heat stress (131), and other kinds of stress (132). Other studies analyze several TEs but not the entire mobilome, such as the case of some overexpressed TE families in the pine of species *Pinus sylvestris* (133). In addition to the focus on specific TEs, a large part of the studies that show TE activation after exposure to stress derive from studies in plants, where there is a high density of TEs in the genome (134). Despite this, the use of large-scale methods to measure the expression, such as microarrays, showed that the number of induced and repressed TEs are almost identical during the heat stress in some species, such as *A. thaliana* (135). Moreover, a recent review (136) shows that TEs are not always activated by stress and that this relationship is quite complex.

The results of this paper focused on several mosquito species bring more content to this debate with the following findings: I) there are stressful situations that do not influence the expression of TEs, II) although there are over-expressed TEs, the number of repressed families was higher than the number of induced ones.

For the first case, we need to consider the different stress types and intensities, and the characteristics of each species and developmental stages. The same type of stress can cause completely opposite responses in distinct species, even if they belong to the same group of animals, or even between different stages of the same animal’s life cycle. As examples of the latter case, here, we observe that diapause-inducing stress affects <2% of TE families of *Ae. albopictus*, but there is an enormous difference between the adult phase and the embryonic phase. In the latter, almost no TE family responds to stress. Even within a specific phase, later age can have a direct impact on the TE response to chemical compounds, as observed during *An. gambiae* exposure to ivermectin. This variation in the TE expression in the face of aging stress appears to be universal, occurring in other dipterans such as *D. melanogaster* to nematodes and rodents (137). Even though aging stress causes similar impacts on such distinct groups of organisms, this does not extend to other types of stress—an elevated level of the same salinity stress, here, produces totally different expression patterns in *An. merus* e *An. coluzzii*. The same stress conditions, such as exposure to malathion, can also result in entirely different responses among *D. melanogaster* lineages, 34 TEs exhibit differential expression in one lineage, whereas in another lineage, malathion does not affect TE expression (138). Thus, it becomes evident that most types of stress, by themselves, do not globally induce the expression of transposable elements in organisms. The impact on TE expression varies among different species and stress intensities.

For the second case, we need to consider that mobilome is diverse, having distinct types of promoters. Moreover, RNA transcription depends on varied factors in the vicinity of a TE locus. In nearly all instances of biotic and abiotic stress to which the mosquito was exposed, there are more repressed than induced TEs. This pattern of expression seems to be more common than expected. A recent review has revealed that several types of stress can modulate the expression of specific *Drosophila* TEs, with repressed TEs outnumbering induced ones among these differentially expressed elements (30). This pattern has also been observed in vertebrates, such as humans, where there are more repressed than induced TEs after exposure to some chemical compounds, such as Simvastatin (139) and short-term treatment with Morphine (140). Classical theories argue that TE induction after stress exposure acts as a force that can generate diversity, consequently contributing to the species’ adaptation to changing environments. However, if most TEs become activated, the high number of transposition events can damage many structures in the genome, contributing to a negative fitness in the host. The prevalence of repressed TEs over induced ones may serve as a stabilizing factor, which allows genetic innovations but reduces the risk of extremely uncontrolled genome damage caused by global mobilome activation.

On the other hand, although the literature often refers to the overexpression of transposon transcripts as a sign of activation and potential transposition, this should be approached with caution. This is because most TE copies are fragmented, implicating their involvement in the formation of long non-coding RNAs (lncRNAs) as previously discussed. In this study, we demonstrate that most transcripts derived from transposons do not encode proteins and, therefore, are likely to contribute as regulatory agents that differentially respond to diverse types of stimuli. Something that has also been observed in *Arabidopsis thaliana*, where TE-lincRNAs are influenced by stress conditions, with the induction or repression response of a specific TE-lincRNA being linked to the type of stress to which the plant has been subjected (141).

Some questions remain as challenges that require complex resolution, for instance: I) Why are some families induced while others are repressed? II) Why are some families more likely to be activated than others? The generation of differentially expressed non-coding transcripts derived from transposable elements in mosquitoes suggests that the expression of some TEs is altered as a cause or consequence of gene expression. If, in the first scenario, variation in TE expression is involved in signaling pathways and regulation for the activation or repression of specific genes, the second scenario poses a new question: does the alteration in TE family expression under stress simply mirror changes in flanking gene expression? Using mainly data from *Ae. aegypti* response to infection, it is clear that the changes in TE expression are not a simple result of gene transcription readthrough. The TE-derived transcripts suffer almost no influence on infection by ZIKV, CHIKV, and DENV viruses in the salivary gland and by CHIKV in the midgut. Although low variation in gene expression has been reported in the latter case (71 differentially expressed gene (DEG) at 1 dpi and 78 DEG at 2 dpi) (53), the variation in TEs is minimal (3 DE in 1 dpi and 0 DE in 2 dpi). In the salivary gland, the number of differentially expressed genes is much higher, with 966 DEGs in CHIKV infection, 396 DEGs in ZIKV infection, and 202 DEGs in DENV infection (55). However, the number of TE DE remains low, with 9 DE in CHIKV infection, 2 DE in ZIKV infection, and 3 DE in DENV infection.

### Relationship of immune response, viral infection, and TE expression

The antiviral response has previously been associated with a strong modulation in the expression of transposable elements. For example, in *D. simulans*, an antiviral response mediated by small RNAs was able to reduce the expression of around 20% of TE families (23). Although both mosquitoes and flies derive from the Diptera clade, the effect of antiviral response seems to be very different (142). We observe that TEs are not strongly modulated by viral infection in *Ae. aegypti*. In fact, in organs experiencing significant viral replication, such as the midgut and salivary gland, the infection causes minimal changes in TE expression. The whole-body analysis of ZIKV infection also showed that less than 3% of TEs respond to infection. However, differently from repression observed in *D. simulans*, the proportion of induced and repressed TEs is almost the same. Although a smaller fraction of mosquito piRNAs map to TEs when compared to species of the genus *Drosophila*. It is possible that a fraction of the repressed TEs, observed in *Ae. aegypti* during this infection with ZIKV, results from the increased production of small RNAs targeting TEs, as a by-product of the production of antiviral defense piRNAs. Despite the absence of evidence for this piRNA antiviral defense by-product in mosquitoes, it has been shown that ZIKV infection can generate piRNAs with no homology to ZIKV, targeting other viral types (37).

We previously observed that some TE-derived lncRNAs function as regulators of genes involved in the antiviral response of *Ae. albopictus* (105). However, it seems that, in *Ae. aegypti*, most transcripts derived from TEs are not involved in immune defense pathways during infection with *B. malayi*, or during ZIKV and CHIKV infection once these infections have minimal effects on the expression of TE families. On the other hand, the first moments of infection with DENV cause a considerable variation in the expression of TEs (196 DETEs at 3 hours post-infection, most of them being repressed), but only 3 genes are DE in the same period (54). This leads us to consider two scenarios. In the first, which is more likely, there is an increase in the number of small RNAs targeting TEs. This increase occurs as a by-product of the overall rise in small RNA levels used to counterattack viral infections. In the second, less likely scenario, changes in TE expression are part of the defense response. In this case, long non-coding RNAs (lncRNAs) would function as regulators of genes involved in the immune cascade. To resolve this puzzle, future investigations with total RNA and small RNA sequencing from the same experimental group are needed to explore whether the production of TE-derived small RNAs is affected by DENV, and other virus, infection. A global analysis must be conducted focusing on comparing the expression of genes, TEs, lncRNAs not involved with TEs and small RNAs.

### Concluding remarks and future directions

The expression pattern of transposable elements is littered with diverging data and remains an openly debated issue. Using extensive transcriptome and proteome datasets, we examined the expression of TEs in mosquitoes. This study reveals that most families of transposable elements are present in mosquitoes, although at significantly lower levels than in many single-copy genes. There is no clear correlation between mobilome expression and the number of TE families. In contrast, Culicinae mosquitoes present a high number of transcript isoforms, driven by the abundance of TE copies in their genomes. However, in most cases, these isoforms result in non-coding RNAs, suggesting a possible regulatory role. The arms race between transposable elements and hosts is reflected in the fact that few families can generate forms of coding RNAs, so some proteins derived from TEs are still found in mosquitoes, presenting a highly tissue-specific profile.

The study also concluded that the expression of TEs is particularly distinct between somatic tissues and germinal tissues. There is, in general, a strong repression of TEs in the ovaries, which seems to be a general rule among animals. The observed overexpression of proteins involved in the piRNA pathway makes the hypothesis of the involvement of piRNAs as effectors of TE repression in the ovary stronger. Finally, considering the diverse types of biotic and abiotic stress, it was shown that the classically accepted argument that TEs are more expressed under stress conditions is not valid for all species/conditions. TEs can be induced by aging, but their responses can vary with the type of stress, intensity, species, or strain subjected to stress. In general, while most TE families in mosquitoes are expressed, only a small fraction are affected by stress stimuli, generally with greater repression of TEs, following a pattern that does not indicate readthrough transcription. These findings highlight the complexity of the interaction, offering new insights into the dynamics between transposable elements and their mosquito hosts.

To conclude, this article outlines issues that deserve consideration in future research. Despite the suggestion of the involvement of the piRNA pathway in the repression of TEs in ovaries, direct validation of the simultaneous expression of piRNAs, mRNAs, and TEs-derived transcripts is crucial to establish the causality of this phenomenon. Regarding the behavior of TEs under stress, although most families keep expression levels, it remains unclear how the reduction observed in some TE superfamilies can influence gene expression more comprehensively. These gaps represent potential areas for future investigation to deepen our understanding of the mechanisms underlying the regulation of transposable elements and their impact on cellular responses.

## DATA AVAILABILITY

Annotation files for TE copies higher than 300 nucleotides, databases used for proteomic analysis, and the list of expressed proteins are available on Figshare https://doi.org/10.6084/m9.figshare.25057466.v1. All RNA-Seq experiments analyzed in the article are available on Sequence Read Archive at NCBI and accession codes are available in Supplementary Tables 1 and 2.

## SUPPLEMENTARY DATA

Supplementary Data are available at NAR online.

## ACKNOWLEDGEMENTS

We thank the Bioinformatics Center of the Aggeu Magalhães Institute and all the researchers responsible for generating the primary data, reanalyzed here from the point of view of transposable elements.

